# Divergent amyloid trajectories distinguish normal aging from Alzheimer’s disease progression

**DOI:** 10.64898/2026.09.08.748727

**Authors:** Mingzhao Tong, Tianchuan Gao, Yurika Upadhyaya, Kwangsik Nho, Shiaofen Fang, Andrew J. Saykin, Jingwen Yan, Alzheimer’s Disease Neuroimaging Initiative

## Abstract

Aging is the greatest risk factor for Alzheimer’s disease (AD), yet how and when AD-related pathological progression diverges from aging remains poorly understood. This distinction is particularly difficult at early stages, when clinically and biomarker-defined populations contain individuals following fundamentally different trajectories. Here, we model AD progression as a deviation from aging using longitudinal amyloid PET and a self-supervised trajectory-learning framework. In ADNI, the learned trajectory revealed a shared early path that bifurcated into an aging branch and an AD-related branch. The two branches showed distinct profiles in amyloid burden, cognitive decline, risk of progression to AD dementia, and genetic risk. Unseen participants from the independent NACC cohort were projected onto the trajectory without retraining, further reproducing key branch-specific biological and genetic patterns. Together, the bifurcating trajectory provides a biologically grounded framework for resolving early-stage heterogeneity and enabling risk stratification beyond binary amyloid status.

## 1. Introduction

Aging is the greatest risk factor for Alzheimer’s disease (AD), yet not all older adults develop AD^1^. Early cohorts therefore include individuals with distinct future trajectories, with some along an aging trajectory and others progressing toward AD. While disease-modifying interventions are widely expected to have the greatest potential during early disease stages^2,3^, determining the direction of progression at this stage remains challenging because current definitions of early stage do not reflect a uniform biological state. In traditional clinical staging, cognitively normal (CN) participants are defined primarily by cognitive performance. However, some CN participants already harbor early amyloid pathology while remaining cognitively unimpaired^3,4^. Others remain cognitively resilient despite accumulating amyloid burden^5–8^.

Biomarker-based frameworks like the ATN have increased the biological specificity of AD diagnosis and staging^9^. Their focus on core AD biomarkers enables disease-related biological changes to be identified and staged before substantial cognitive impairment becomes apparent. However, within these frameworks, biomarker status is often interpreted categorically, such as amyloid-positive or amyloid-negative, and this binary distinction does not fully resolve heterogeneity in underlying disease progression. Substantial variability in clinical and cognitive outcomes has long been recognized within preclinical AD populations^10,11^.

Amyloid-negative populations are generally enriched for individuals who are younger or at an earlier position along the amyloid-accumulation continuum^12^, making their future trajectories particularly uncertain. This broad biomarker category can therefore encompass individuals who remain at low amyloid levels with stable cognition as well as those undergoing early amyloid accumulation that has not yet crossed the conventional positivity threshold. In the amyloid-negative LEARN cohort, 73.8% of cognitively unimpaired participants remained clinically stable over approximately 4.5 years, whereas 26.2% showed sustained progression to cognitive impairment^13^. In line with this, subthreshold amyloid levels among amyloid-negative individuals have also been associated with subsequent tau deposition and cognitive decline^14,15^, providing strong evidence that some amyloid-negative individuals may already be undergoing early AD-related pathological change. Substantial variability also remains after individuals become amyloid positive. Among amyloid-positive cognitively normal participants in the AIBL cohort, approximately 25% progressed to MCI or dementia over a mean follow-up of 5.3 years, with progression risk ranging from 12% among those with moderately elevated amyloid to 50% among those with very high amyloid levels^16^. Taken together, individuals sharing the same clinical status and binary amyloid classification can nevertheless follow markedly different future trajectories. A framework that characterizes these diverging trajectories may therefore provide greater biological resolution than binary amyloid status alone and improve the biological characterization of early AD^17^.

To address this gap, we model AD progression as a deviation from aging using longitudinal amyloid PET data from ADNI. We identify a bifurcating trajectory in which participants initially share a common early path before diverging into an aging branch and an AD-related branch.

These branches show distinct patterns of amyloid accumulation, cognitive decline, clinical progression risk, and APOE-related genetic risk. We further demonstrate that the learned trajectory generalizes to the independent NACC cohort, where key branch-specific biological and genetic patterns are reproduced without retraining. This data-driven staging framework moves beyond binary amyloid classification to provide a more refined and biologically grounded representation of AD progression, with the potential to improve risk stratification in both research and prevention-oriented settings.

## 2. Results

### 2.1. Amyloid trajectories bifurcate into normal aging and AD-related paths

Applied to longitudinal amyloid PET data from ADNI, the learned embedding revealed a clear bifurcating structure in the landscape of amyloid accumulation, comprising a shared preclinical trunk and two diverging branches (Fig. 1a). The shared segment preceding the bifurcation was occupied by participants spanning all diagnostic groups, indicating that it captures an early biological stage at which different aging trajectories have not yet diverged. Beyond the bifurcation, branch composition showed a consistent pattern across the multi-visit training and test cohorts. Participants with stable AD or progressive MCI (pMCI; MCI converting to AD during follow-up) were predominantly localized along one branch, whereas stable cognitively normal participants and stable MCI cases (sMCI) were distributed across both branches. Based on the relative enrichment of AD and pMCI cases along one branch and CN participants along the other, the two branches are hereafter referred to as the AD branch and the aging branch. The same pattern was observed in the single-visit group, where AD cases predominantly aligned with the AD branch, while CN and MCI participants were distributed across both branches and the shared trunk.

**Figure. 1:**
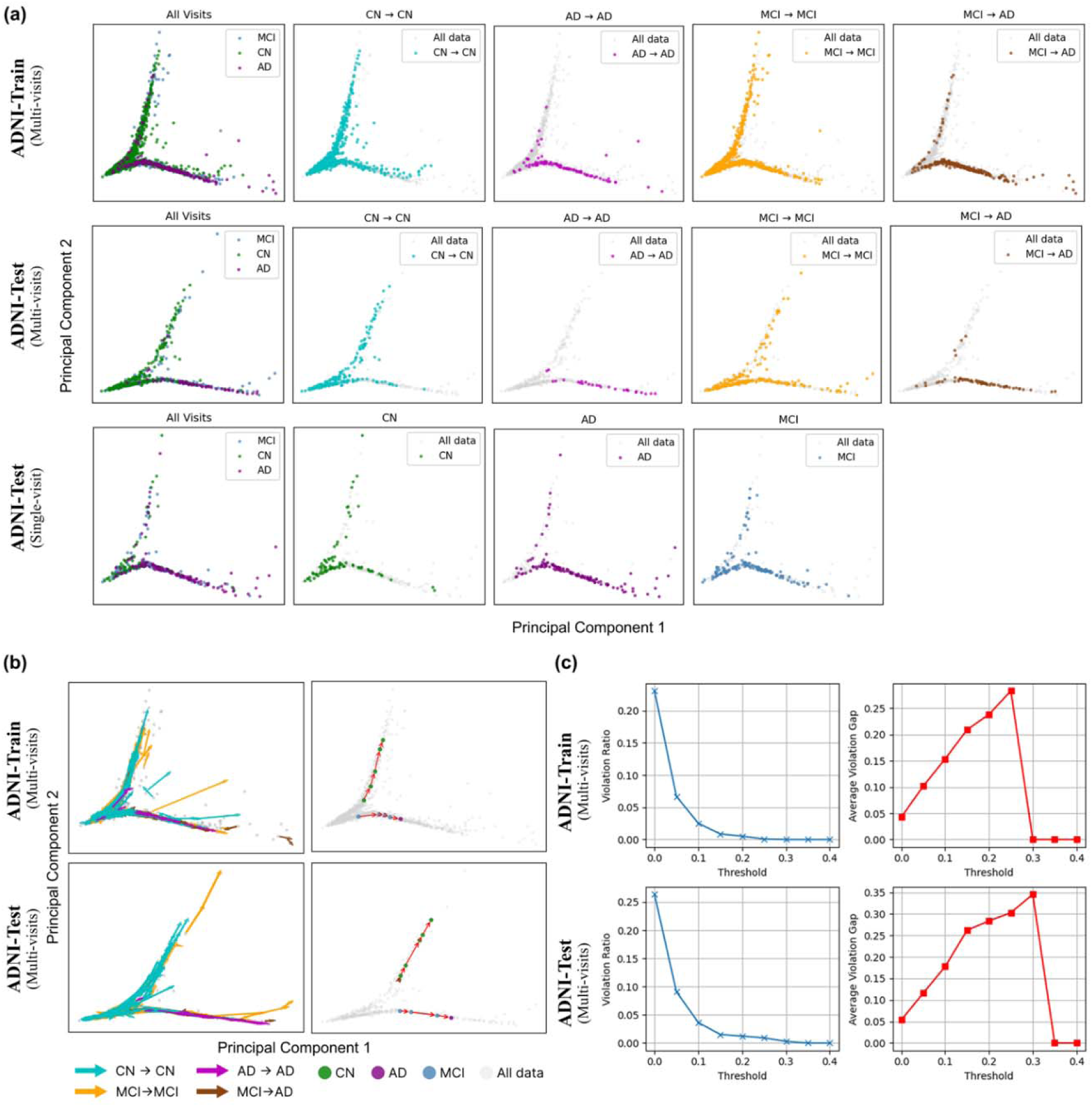
Branched trajectory and temporal consistency in ADNI. (a) Two-dimensional branched trajectories of ADNI regional SUVR data across the training and test cohorts. Three panels are for ADNI training set, ADNI test set with multiple visits and ADNI test set with one single visit respectively. (b) Longitudinal progression in multi-visit training and test sets, with subject trajectories colored by diagnostic transition and representative serial visits connected along inferred branches. (c) Temporal consistency across follow-up visits. Tolerance thresholds are set to ignore the minor reversal of pseudotime between consecutive follow-up visits. Lower values in violation ratio and violation gap indicate greater temporal consistency.

We further quantified the diagnostic separation and longitudinal stability of the branches. Branch membership was used to differentiate amyloid-negative CN participants from AD cases, excluding participants on the shared trunk due to its diagnostically mixed composition (Fig. S1, Table S1). We restricted the CN group to amyloid negative participants with Centiloid < 10 to reduce within-group heterogeneity and provide a cleaner contrast with AD cases. Classification performance was consistently strong across training and held-out multi-visit test groups (accuracy ≥ 0.89, F1 ≥ 0.91) and remained strong in the single-visit test group, indicating that the bifurcation structure learned from longitudinal data generalizes to cross-sectional observations. Longitudinal follow-up visits also generally progressed forward along the inferred branches (Fig. 1b,c). Each individual was assigned a continuous staging score (ranging from 0 to 1), reflecting their relative distance to the trajectory root. Fewer than 5% of consecutive visit pairs showed reversed staging score at a tolerance threshold of 0.1 in both training and test cohorts (e.g., minor reversal in staging score (<0.1) was ignored). Branch assignments were similarly stable over follow-up visits. During the follow-up period, branch switching was rare, occurring in only 16 of 551 participants in the training set and 7 of 154 in the test set. Furthermore, almost all transitions took place at the very beginning of bifurcation, a point at which the two branches had not yet shown clear divergence (Fig. S2). Together, these results demonstrate a diagnostically coherent, temporally ordered, and stable bifurcating representation of amyloid progression.

### 2.2. AD-related and aging branches show distinct amyloid, cognitive and age profiles

We examined associations between branch-specific staging score and regional amyloid burden, cognitive performance, and age across three ADNI subsets: multi-visit training set, held-out multi-visit test set and held-out single visit test set (Fig. 2). Along the AD branch, regional amyloid burden increased progressively with staging score, with a steeper increase after the bifurcation point, and this pattern was consistent across all three subsets. Higher staging scores in AD branch were also associated with poorer memory (r = −0.44 to −0.49, p < 0.0001) and executive function (r = −0.38 to −0.41, p < 0.0001). These associations remained significant in the single-visit subset (memory: r = −0.49, p < 0.0001; executive function: r = −0.38, p < 0.0001), whereas age showed only a modest association with staging score on the AD branch (r = 0.19– 0.23, p < 0.001).

**Figure. 2:**
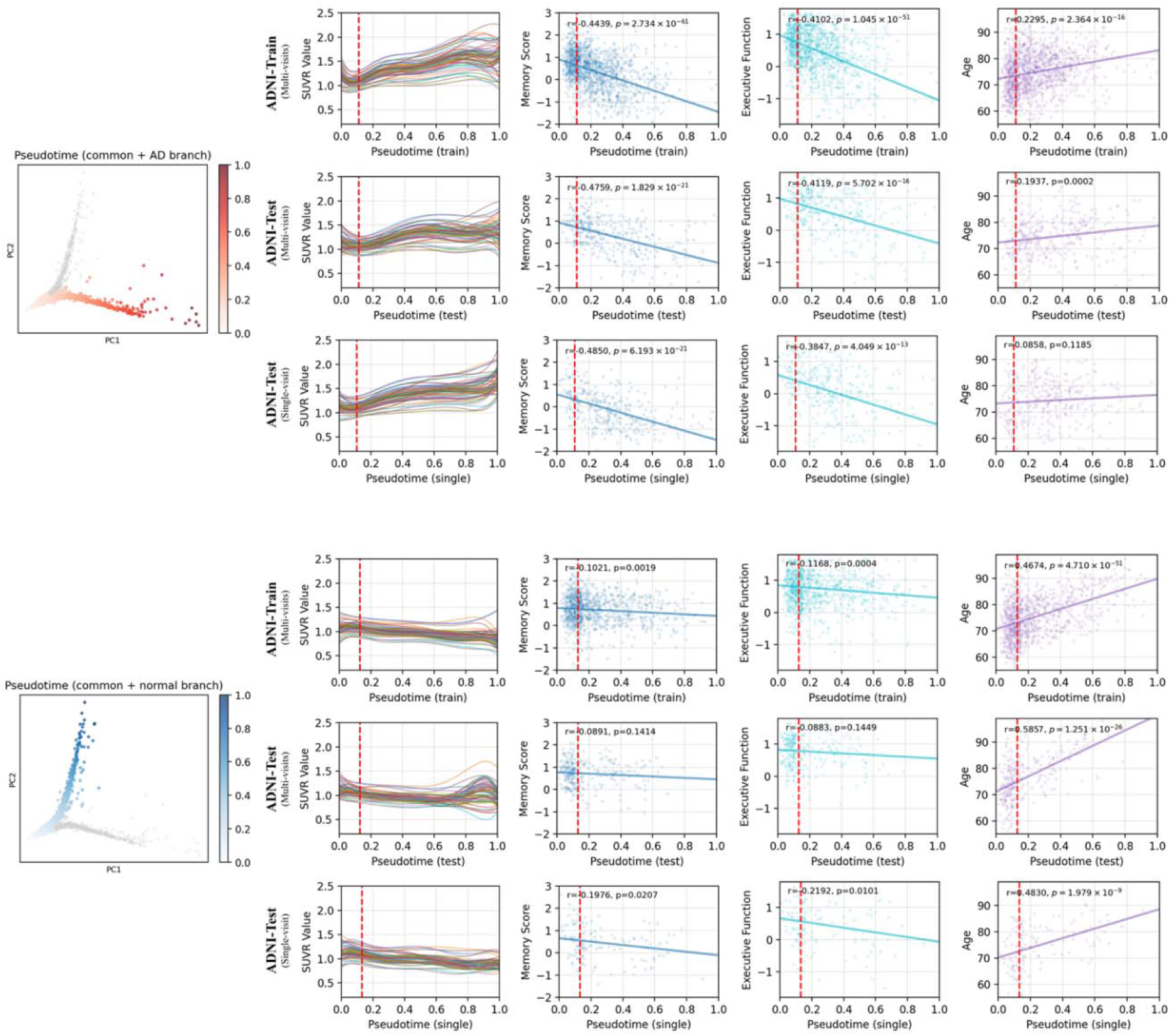
Association of branch-specific staging scores with amyloid burden, cognition, and age in ADNI. Top panels show the AD-related branch; bottom panels show the aging branch, each including the shared trunk. Staging score is the normalized distance from the trunk root along each branch and is mapped onto the embeddings at left. Regional amyloid SUVR trajectories are shown for the training, held-out test with multiple follow-up visits, and test participants with one single visit (marked as single); dashed lines mark the bifurcation point. Scatter plots show associations with memory (PHC_MEM), executive function (PHC_EXF), and age with linear fits. Pearson correlations (r) and p values are reported.

The aging branch showed a distinct profile. Regional amyloid burden remained relatively stable across staging score, consistent with removal of the linear age effect from regional SUVR before embedding, and associations with cognition were substantially weaker than on the AD branch. Memory correlations were weak or non-significant in the training set (r = −0.10, p = 0.0019) and in the held-out test subsets (multi-visit test: r = −0.09, p = 0.1414; single visit test: r = −0.20, p = 0.0207), with similar patterns for executive function (train: r = −0.12, p = 0.0004; test: r = −0.09, p = 0.1449; single visit: r = −0.22, p = 0.0101). In contrast, age was strongly associated with staging score (train: r = 0.47, p < 0.0001; test: r = 0.59, p < 0.0001; single visit: r = 0.48, p < 0.0001). Thus, the two branches showed distinct profiles. The AD branch primarily tracked amyloid accumulation and cognitive decline, whereas the aging branch was more strongly associated with chronological age.

### 2.3. Branch membership is associated with genetic variation at the APOE locus

Genome-wide association analysis of branch membership identified a prominent signal on chromosome 19 spanning the *NECTIN2–TOMM40–APOE–APOC1* region and reaching genome-wide significance (p < 5 × 10^-8^; Fig. 3). The signal was observed in the full sample and remained evident in the CN/MCI subset, indicating that the association was not driven solely by clinically diagnosed AD cases. Consistent with the established role of *APOE* in AD risk, the rs429358-C risk allele was more frequent among participants on the AD branch than on the aging branch in both the full sample and CN/MCI subset. These results show that the imaging-derived bifurcation aligns with established *APOE*-related genetic risk, including at early clinical stages.

**Figure. 3:**
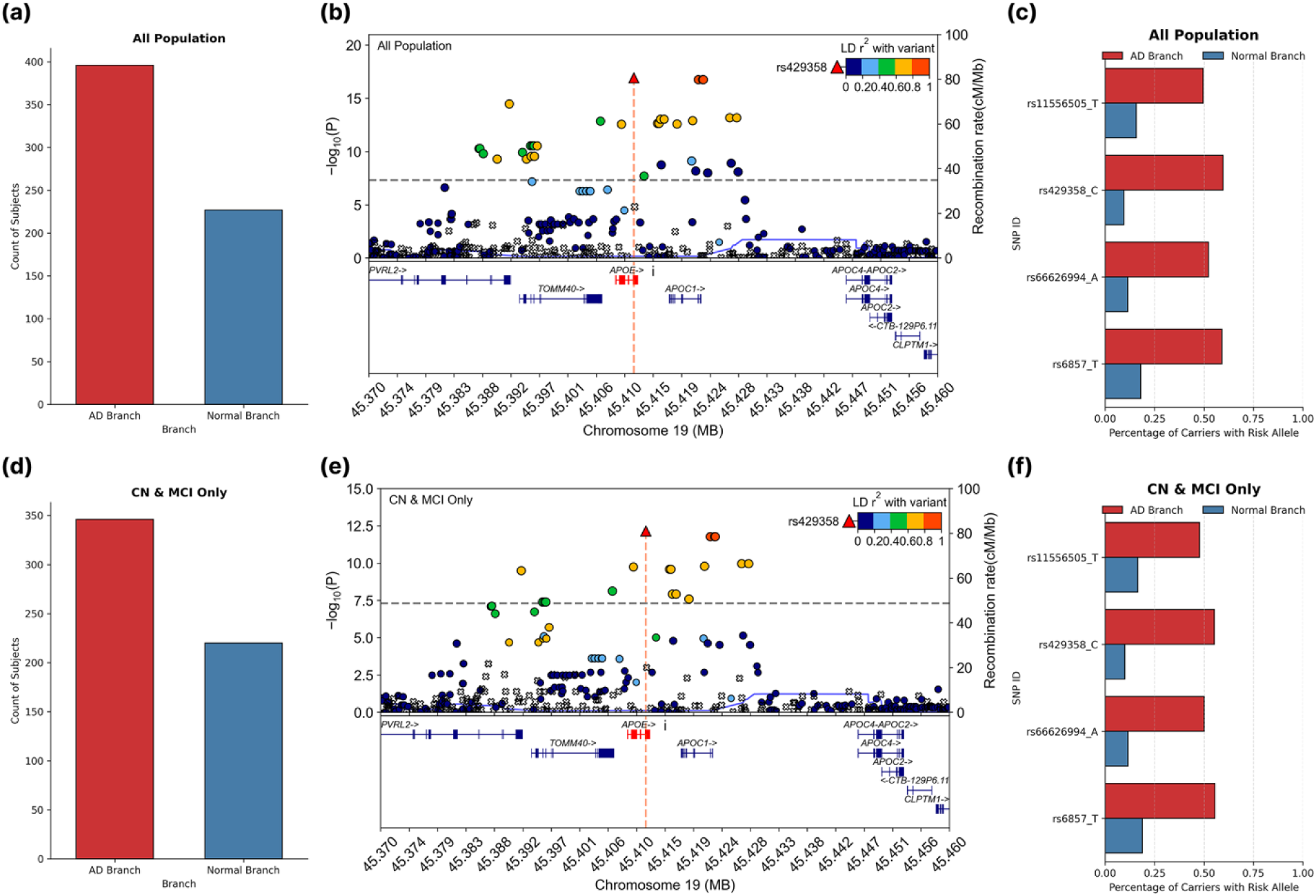
Branch-associated genetic variation at the *APOE* locus in ADNI. Left: sample sizes for the AD-related and aging branches in the full cohort (a) and CN/MCI subset (d). Center, top genetic variants associated with branch membership in all ADNI samples (b) and CN/MCI subset (e); dashed lines indicate genome-wide significance (P < 5 × 10^-8^). Right: frequencies of representative risk alleles in the full ADNI cohort (c) and CN/MCI subset (f). Red denotes AD-related branch and blue denotes aging branch.

### 2.4. Branch membership predicts longitudinal cognitive decline and AD conversion risk

Among participants with CN and MCI at baseline, their branch membership was strongly associated with subsequent cognitive decline and clinical progression. In the ADNI training set, participants assigned to the AD-related branch showed significantly faster decline in both memory (p = 2.52 × 10^-22^; Fig. 4a) and executive function (p = 1.03 × 10^-13^; Fig. 4b) compared to those on the aging branch. Survival analysis further demonstrated that CN/MCI participants on the AD-branch had a substantially higher risk of conversion to AD dementia over follow-up (p = 7.44 × 10^-6^; Fig. 4c). These findings were replicated in the held-out test set, where AD branch participants again showed significantly faster memory decline (p = 5.74 × 10^-5^; Fig. 4d), faster executive function decline (p = 2.54 × 10^-3^; Fig. 4e), and higher conversion risk (p = 9.29 × 10^-3^; Fig. 4f). These results indicate that branch membership derived solely from amyloid PET data captured clinically meaningful differences in future cognitive and diagnostic trajectories among participants sharing the same broad baseline clinical status.

**Figure. 4:**
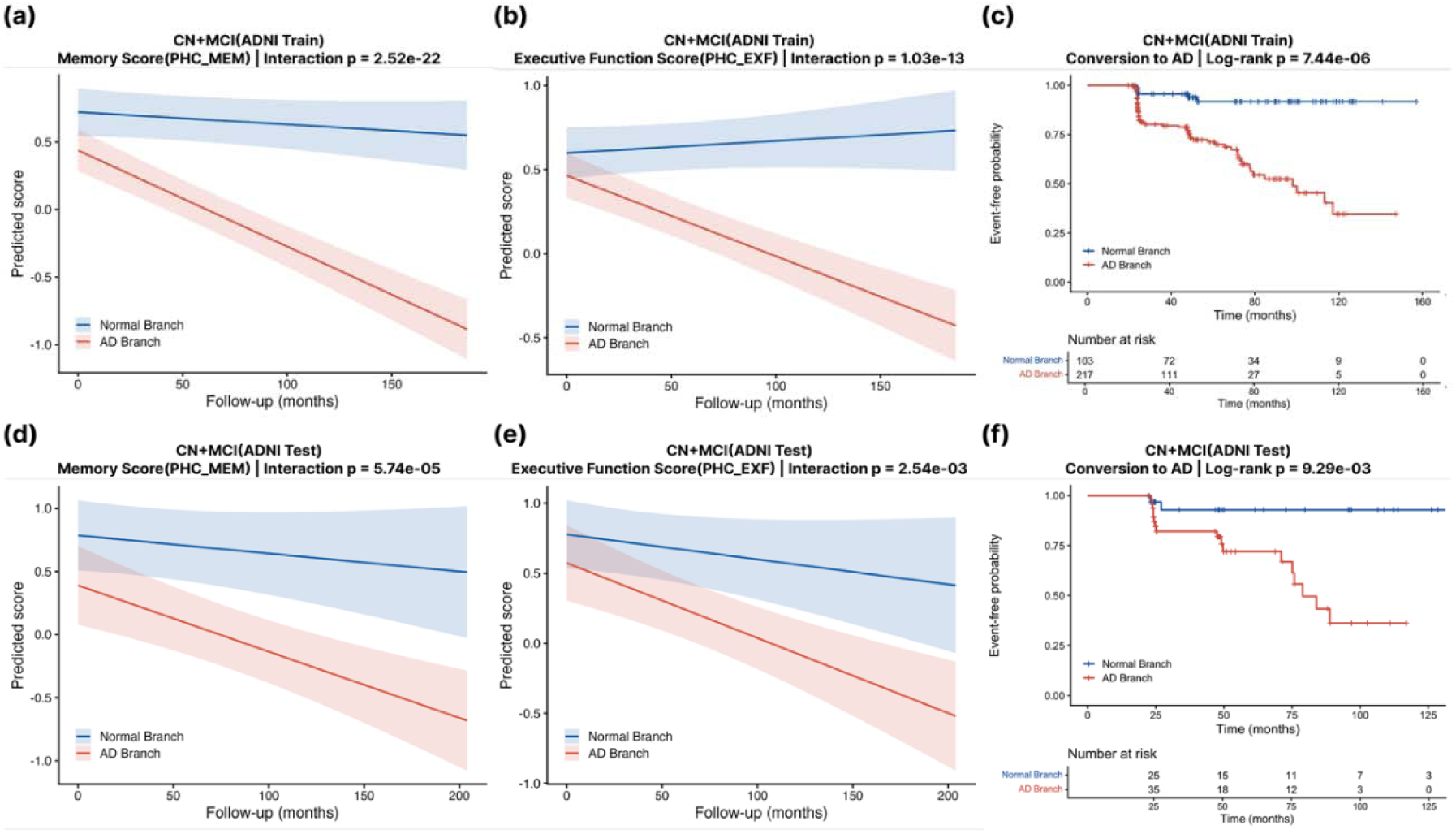
Branch-specific longitudinal cognitive decline and AD conversion risk in ADNI. CN and MCI participants were assigned to the AD-related branch (red) or aging (blue) branch based on their projected location. In the ADNI training set, CN/MCIs located on the AD branch showed faster decline in memory score (a) and cognitive function (b), and higher risk of conversion to AD (c). Similar results were also observed in the held-out test set (d–f).

### 2.5. The bifurcating trajectory generalizes to an independent external cohort

To assess generalizability, the trained trajectory framework was applied without retraining to the independent NACC cohort, for which harmonized amyloid PET data were available only from single visits. Projection of NACC participants onto the ADNI-trained trajectory reproduced the branch-specific diagnostic pattern observed in ADNI, with AD cases concentrated along the AD branch and CN/MCI participants distributed across both branches and the shared trunk (Fig. 5a).

**Figure. 5:**
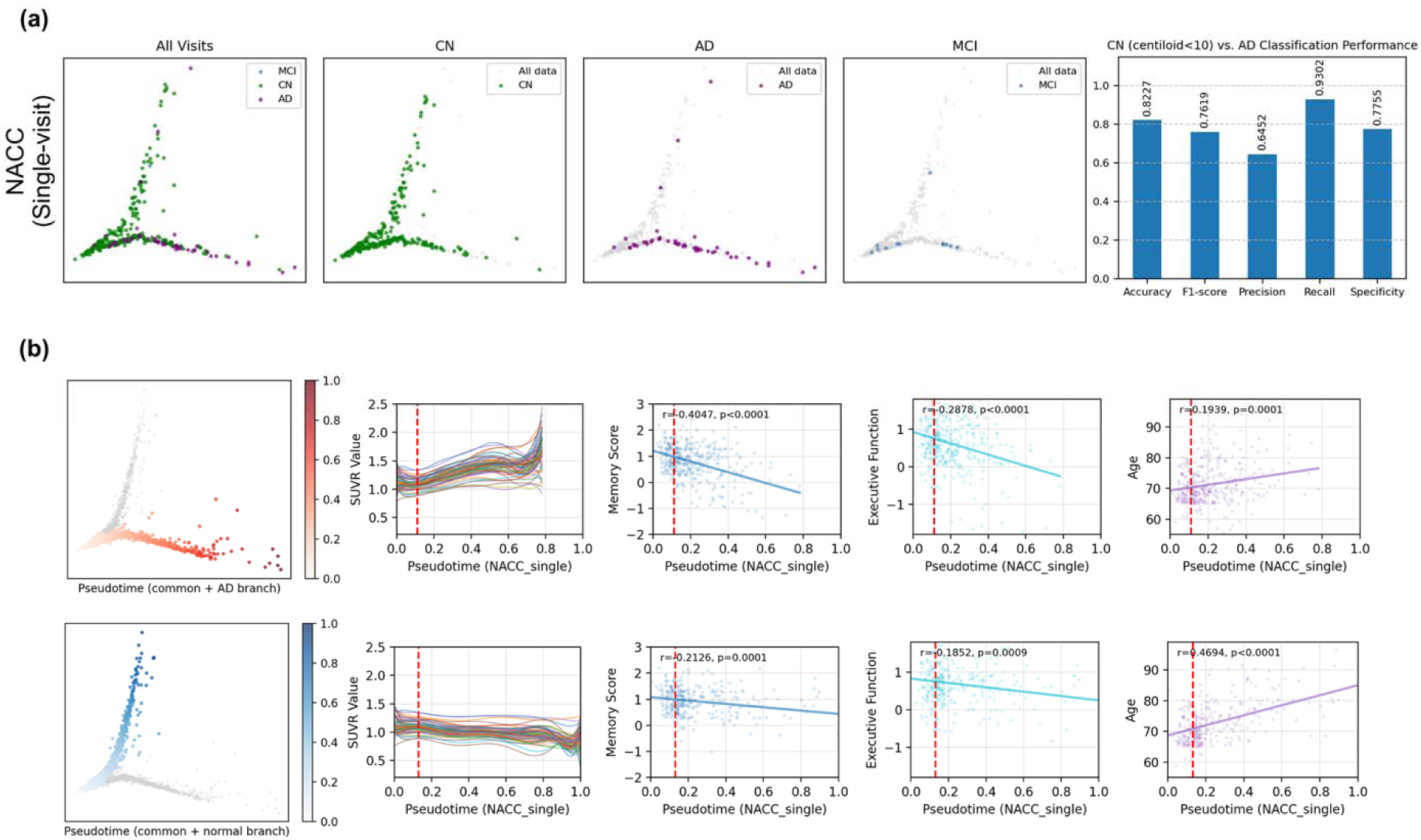
External validations of the learned trajectory framework in NACC. (a) NACC embedding by diagnosis and branch-based classification of AD versus amyloid-negative CN participants (Centiloid <10). (b) Associations of branch-specific staging scores with regional amyloid SUVR, memory, executive function, and age for the AD-related branch (top) and aging branch (bottom). Dashed lines mark the bifurcation point; Pearson correlations and two-sided p values are shown.

Branch-based classification of amyloid negative CN participants versus AD cases remained robust in NACC, achieving an accuracy of 0.823, F1-score of 0.762, precision of 0.645, recall of 0.930, and specificity of 0.776 (Fig. 5a). The higher recall relative to precision indicates high sensitivity for AD cases, with lower specificity of AD-branch assignment across the clinically heterogeneous NACC cohort. Branch-specific biological profiles were also reproduced in NACC (Fig. 5b). On the AD branch, higher staging scores were associated with greater amyloid burden and poorer memory (r = −0.40, p < 0.0001) and executive function (r = −0.28, p < 0.0001), with only a modest association with age (r = 0.19, p = 0.0001). On the aging branch, associations with memory (r = −0.21, p = 0.0002) and executive function (r = −0.19, p = 0.0007) were weaker, whereas age showed a much stronger association with staging score (r = 0.47, p < 0.0001).

*APOE*-related differentiation between branches was also reproduced in NACC. Among CN/MCI participants, *APOE* ε4 carriers were more likely to be assigned to the AD branch than to the aging branch (χ² = 13.06, p = 3.0 × 10^-4^). Longitudinal cognitive decline and AD conversion analyses were not feasible because only one diagnostic conversion was observed among CN/MCI participants during follow-up. Together, the diagnostic, biological, and *APOE*-related patterns observed in NACC support the generalizability of the ADNI-derived trajectory to an independent cohort.

### 2.6. AD-branch pseudotime captures early amyloid progression and predicts cognitive decline

Progression along the AD branch (including shared trunk) is quantified using pseudotime (PT_AD), calculated as the relative position to the trunk root (ranging [0,1]). Across all participants, PT_AD showed a consistently lower pairwise violation ratio than global amyloid SUVR across tolerance threshold (Fig. 6a), indicating more coherent longitudinal ordering. Both global amyloid SUVR and PT_AD were higher in MCI relative to CN participants, but group separation was substantially stronger for PT_AD (p = 8.816 × 10^−9^) than for global amyloid SUVR (p = 0.005334) (Fig. 6b). The bifurcation point marking clear divergence toward the AD-related trajectory was observed at approximately PT_AD = 0.22 (Figure 6c). Among all AD-branch participants before this point (PT_AD ≤0.22), bilateral precuneus amyloid showed the strongest association with PT_AD, with the left and right precuneus yielding the largest explained variance and strongest statistical significance among regional SUVR measures (Fig. 6d). Finally, to evaluate the prognostic value of PT_AD within a clinically relevant range of amyloid burden, we focused on CN/MCI participants with Centiloid values of 20–80. This range spans intermediate to elevated amyloid levels and captures the early amyloid stages often targeted by prevention trials^18^. Within this subgroup, participants with PT_AD >= 0.22 showed significantly faster memory decline than those with PT_AD <0.22 in both the training cohort (N = 197, p = 4.91 × 10^-7^) and held-out test cohort (N = 57, p = 0.00551; Fig. 6e). These findings support the relevance of AD-branch pseudotime for characterizing early amyloid progression and subsequent cognitive decline.

**Figure. 6:**
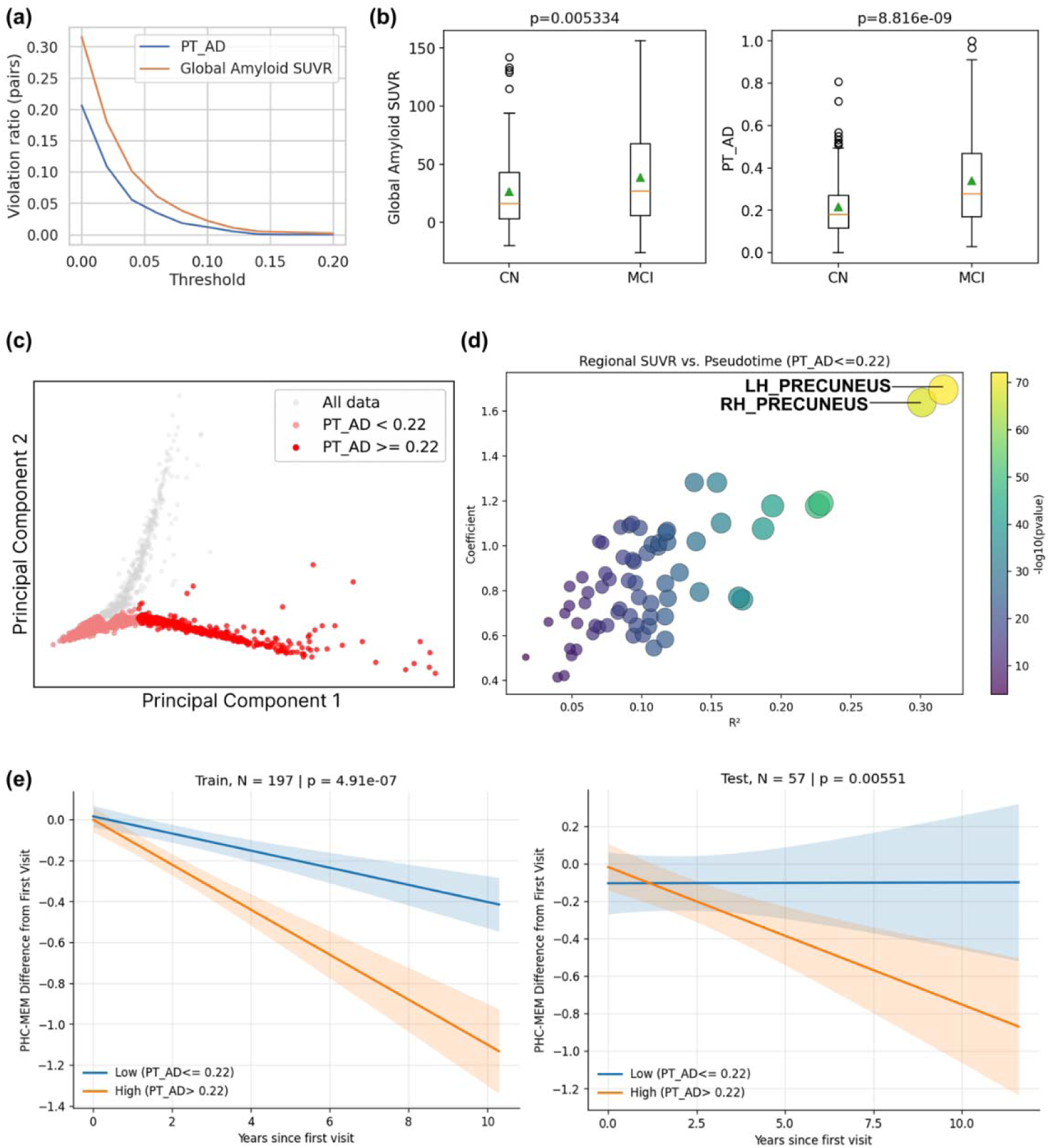
Evaluation of AD-branch progression. (a) Comparison of violation ratios for AD-branch staging score (PT_AD) and global amyloid SUVR. (b) Distributions of global amyloid SUVR and PT_AD in CN and MCI groups. (c) AD-branch cohort stratified at PT_AD = 0.22. (d) Associations between PT_AD and individual regional amyloid SUVR among participants with PT_AD ≤ 0.22. (e) Longitudinal memory decline of CN/MCI subjects with intermediate to elevated amyloid levels. Subjects were stratified by the bifurcating point (PT_AD = 0.22). Left: ADNI training set. Right: ADNI test set.

## 3. Discussion

Aging and AD related pathological change coexist in older adults, yet their early trajectories are difficult to distinguish using clinical diagnosis or categorical biomarker status alone. Here, longitudinal amyloid PET revealed a shared early path that bifurcated into an aging branch and an AD related branch. This structure provides a data driven perspective on the longstanding question of whether AD represents accelerated aging or a distinct pathological process. Aging and AD share many biological and cognitive changes, which has supported the view that AD may reflect an acceleration of processes occurring during aging^19^. However, qualitative differences between aging and AD argue against explaining AD simply as faster aging^20^. Our finding of a shared early path followed by bifurcation supports an intermediate view. Aging related and AD related amyloid trajectories may initially overlap but become progressively distinct as AD related pathology emerges. This reframing suggests that the transition toward AD related progression can’t be fully explained by accelerated aging but instead may reflect a gradual divergence that unfolds over years^21,22^.

A key insight from the trajectory analysis is that the shared preclinical trunk represents a stage in which progression direction remains unresolved. It is occupied by participants across diagnostic groups, in whom aging is ongoing but the direction of amyloid progression is not certain yet.

Current biomarker frameworks, including the revised AT(N) criteria, provide limited resolution of this ambiguity^9^. The bifurcation identified by our model may represent the stage at which aging related and AD related progression begins to diverge. Some participants continue along a stable aging path, while others transition toward an AD related amyloid accumulation trajectory. The stable branch membership across follow-up visits after the major bifurcation suggests that progression patterns become more clearly established once the trajectories separate. In contrast, individuals near the bifurcation or within the shared trunk remain less clearly differentiated in the current framework, and their subsequent progression direction cannot yet be reliably resolved.

The biological factors that drive this observed divergence remain uncertain and may include differences in amyloid accumulation, regional spread, cognitive reserve, and genetic vulnerability^5,8,23^. Resolving this uncertainty will likely require the integration of multimodal biomarkers into future trajectory frameworks^24^.

The branch specific associations with amyloid, cognition, and age further clarify the biological meaning of this divergence. Progression along the AD branch was associated with increasing amyloid burden and worsening memory and executive function, while its association with chronological age was modest. In contrast, progression along the aging branch showed much weaker relationships with amyloid and cognition but a stronger association with age. This asymmetry suggests that progression along the two branches is organized by different dominant sources of variation. Along the AD-related branch, staging score appears to track increasing pathological burden together with its clinical consequences, whereas along the aging branch, pseudotime is more closely aligned with chronological aging in the relative absence of comparable amyloid accumulation or cognitive decline. Clinical and genetic findings provide additional support for this interpretation. Among CN and MCI participants, AD branch membership was associated with faster cognitive decline and greater risk of conversion to AD dementia. Branch membership was also associated with genetic variation across the *APOE* region, even within CN and MCI participants. This alignment with established AD genetic risk further speaks to the biological validity of the trajectory^25–27^. Together, these findings suggest that once progression direction becomes clearly differentiated, branch membership is associated with biologically and clinically meaningful differences at the group level.

The early portion of the AD-related trajectory provides a closer view of progression before the two paths become clearly separated. Because age effects were removed from regional amyloid measures before trajectory learning, progression in the learned space reflects regional variation beyond what is expected from age. Before the clear bifurcation, AD-branch staging score was most strongly associated with bilateral precuneus amyloid. The precuneus and adjacent posterior cingulate cortex have been frequently implicated in early amyloid accumulation^5^. The emergence of the precuneus through its association with trajectory position supports the biological relevance of the progression captured along this early AD related path.

Thus, even before the trajectories become clearly separated, movement along the inferred AD related trajectory reflects meaningful regional amyloid variation. The clinical relevance of location relative to the major bifurcation was also supported among CN and MCI participants with intermediate to elevated amyloid burden. Participants beyond the approximate bifurcation showed faster subsequent memory decline than those earlier on the trajectory. Future studies should determine whether higher staging scores within the resolved AD-related branch are associated with progressively faster cognitive decline or greater clinical risk in larger longitudinal and prevention cohorts.

Looking forward, an important challenge is determining how well these trajectory patterns generalize across the broader older adult population. Aging and AD-related progression are shaped by genetic, lifestyle and many other factors that vary across individuals and populations^8,23,28^. Future studies should therefore evaluate these trajectories in more diverse cohorts, including groups underrepresented in current AD studies, in whom the relationship between aging biology and AD-related pathology may differ^29^. Translation to clinical and research settings will also require validation across imaging platforms, tracers, and data collection settings, together with prospective evidence that trajectory-based staging provides meaningful value for prevention-oriented studies and interventions. Together, these efforts may support a more continuous and biologically grounded understanding of how aging and AD related pathology diverge over life course.

## 4. Methods

### 4.1. Datasets

In this study, we utilized data from two independent cohorts: the Alzheimer’s Disease Neuroimaging Initiative (ADNI) and the National Alzheimer’s Coordinating Center (NACC)^30,31^. Both datasets were accessed in a harmonized format through the ADSP Phenotype Harmonization Consortium (PHC), which standardizes variables across cohorts to facilitate cross-dataset analyses^32^. At the time of analysis, harmonized longitudinal data were not yet fully available across all cohorts. Therefore, we used both longitudinal and baseline-only data from ADNI, whereas analyses in NACC were restricted to baseline observations. At each visit, clinical diagnosis was categorized as cognitively normal (CN), mild cognitive impairment (MCI), or Alzheimer’s disease (AD). Cognitive performance was assessed using PHC-harmonized measures, including PHC_MEM for memory and PHC_EXF for executive function. For both datasets, only white and non-Hispanic participants were included for the analysis.

Amyloid burden was quantified using standardized uptake value ratio (SUVR) derived from amyloid PET imaging^33–35^. For model development, we used volume-weighted mean SUVR values across 68 bilateral cortical regions of interest, capturing regional amyloid distribution patterns. These regional SUVR features were used as input to the unsupervised trajectory model. SUVR measures from all cohorts were harmonized through the PHC pipeline to ensure comparability across imaging protocols and sites, including normalization procedures, reference regions, and scaling methods. To further ensure consistency in regional SUVR measurements, only imaging scans acquired using florbetapir (FBP) tracers were included in the analysis.

#### 4.1.1. ADNI

Data used in the preparation of this article were obtained from the Alzheimer’s Disease Neuroimaging Initiative (ADNI) database (adni.loni.usc.edu). The ADNI was launched in 2003 as a public-private partnership, led by Principal Investigator Michael W. Weiner, MD. The primary goal of ADNI has been to test whether serial magnetic resonance imaging (MRI), positron emission tomography (PET), other biological markers, and clinical and neuropsychological assessment can be combined to measure the progression of mild cognitive impairment (MCI) and early Alzheimer’s disease (AD).

In this study, a total of 1066 participants from ADNI were included. At baseline, these comprised 384 CN, 498 MCI, and 184 AD patients. These participants contributed a total of 2543 visits, with an average of 2.39 visits per subject (range:1–7). Among all participants, 361 participants had only a single visit, including 85 CN, 149 MCI, and 127 AD patients. The remaining 705 participants had longitudinal follow-up data (≥2 visits). Within the longitudinal subset, disease progression was observed in a subset of participants. Specifically, 86 participants with MCI at baseline converted to AD, and 55 participants with CN at baseline converted to MCI or AD during follow-up. The clinical diagnoses of the remaining participants remained stable across visits. These longitudinal data provide a valuable cohort for characterizing disease progression and for evaluating heterogeneity among early-stage participants within the trajectory framework. Supplementary Figure S3 provides an overview of subject distribution and diagnostic categories within the ADNI cohort. Detailed demographic information of ADNI participants is shown in Supplementary Table. S2.

#### 4.1.2. NACC

The National Alzheimer’s Coordinating Center (NACC) aggregates longitudinal standardized clinical, cognitive, neuropathology and biomarker data from more than 40 Alzheimer’s Disease Research Centers (ADRCs) in the United States, from the mid-1980s onwards and with unified data collection from 2005 via its Uniform Data Set (UDS)^36^.

In this study, a total of 321 participants from NACC were included. At baseline, these comprised 260 cognitive normal (CN), 12 mild cognitive impairment (MCI), and 49 Alzheimer’s disease (AD) participants. Due to the availability of harmonized data, amyloid analyses in NACC were restricted to baseline observations. Longitudinal cognitive and diagnostic information were available for a subset of participants, but very limited conversions were observed. Specifically, only 1 CN participants converted to MCI or AD, and none of MCI participants converted to AD during follow-up. The majority of participants remained clinically stable over the observation period. The relatively low number of conversion events and the heterogeneity of the NACC cohort limit statistical power for detecting longitudinal prognostic effects. Accordingly, analyses in NACC were primarily focused on cross-sectional validation. Detailed demographic information of NACC participants is shown in Supplementary Table. S2.

### 4.2. Model

We developed a three-step pipeline to construct a longitudinal trajectory representation of amyloid dynamics, consisting of (1) covariate adjustment, (2) longitudinal embedding, and (3) 2D trajectory learning.

#### Step 1: covariate adjustment

To reduce potential confounding effects of age and sex, we first preadjusted all regional SUVR values using a linear regression model estimated from baseline cognitively normal (CN) participants in the ADNI training set. This helps center all regional values relative to the CN population while preserving disease-related variation. This linear model was saved and subsequently applied to all test and external datasets.

#### Step 2: longitudinal embedding

We adapted the longitudinal neighborhood embedding (LNE) framework^37^ which was originally developed for high-dimensional 3D imaging data. We retrained a low-dimensional version for regional amyloid SUVR values. We employed an autoencoder architecture with symmetric encoder–decoder structure. The encoder maps each visit’s 68-dimensional regional SUVR vector to a lower-dimensional latent representation, while the decoder reconstructs the input from the latent space. Longitudinal information was incorporated by constructing ordered visit pairs for each subject. For each subject with multiple visits, all possible pairs (X(1),X(2))were generated such that corresponds to an earlier visit and to a later visit. The inputs of LNE model are these pair-wise longitudinal data, where both visits are independently encoded and reconstructed. The training objective combines a reconstruction loss and a directional loss:

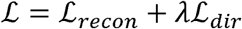

The reconstruction loss ensures fidelity of the input features:

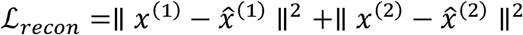

The directional loss constrains the latent difference 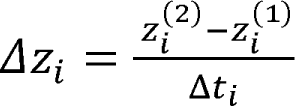 to align with temporal progression of nearby participants. A local directional vector is estimated from neighborhood structure in the latent space:

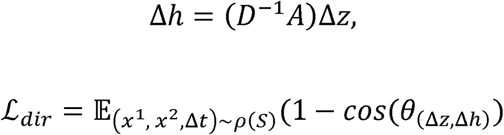

where is the adjacency matrix and is the corresponding degree matrix. This encourages latent trajectories to follow consistent progression patterns across participants .

#### Step 3: progression trajectory learning

The longitudinal embedding generated from the encoder of LNE model will be further reduced to 2D using principal component analysis (PCA)^38^. PCA parameters were learned exclusively from the training data and then applied to test and external datasets to ensure consistent projection. The resulting two-dimensional embedding provides a continuous trajectory representation, from which branching structure and pseudotime can be inferred.

All components of this pipeline were pre-trained and saved after training, including the covariate adjustment model, encoder in longitudinal embedding, and PCA transformation. These pre-trained components were applied to all held-out test data and external datasets, enabling consistent projection of new samples onto the learned trajectory without retraining.

### 4.3. Branch identification

We employed an iterative process to detect the branches in the learned 2D progression trajectory. We iteratively cluster samples using -means^39^ with decreasing cluster numbers. For each candidate , cluster centroids are extracted and used to estimate the topology of the trajectory. We construct a nearest-neighbor among centroids, followed by a minimum spanning tree (MST)^40^, to ensure a connected and acyclic structure. Terminal nodes (degree = 1) in the MST define possible trajectory endpoints, and the first observed terminal node is selected as the root. Branch points are defined as MST nodes with ≥3. For each branch point, we traverse outgoing edges via breadth-first search and record continuous paths until another branch point or terminal node is reached, yielding discrete trajectory branches. We experimented with multiple cluster numbers (range: 8-50) and used the one that consistently produced a stable bifurcation pattern. The final trajectory was characterized by a tree structure consisting of a shared trunk and two diverging branches, corresponding to aging and AD-related progression.

### 4.4. Pseudotime Estimation

Pseudotime was estimated separately for the two branches of the trajectory, indicating the relative distance of each visit to the root in the trunk. For each branch (together with shared trunk), we fitted a smooth spline curve through the ordered points along that trajectory. Each visit along the trajectory was projected onto a densely sampled version of the spline, and its pseudotime was defined as its relative position along the fitted curve, measured as the normalized arc length from the root to that visit. Pseudotime of 0 indicates the root or starting point of the trajectory, and 1 means the end of the branch. Intuitively, pseudotime represents how far the amyloid of a subject has progressed along a given trajectory, with values increasing from early to late stages within each branch. The fitted spline models for both branches were fixed after training and applied to all test and external datasets, ensuring consistent projection and comparable pseudotime across datasets.

### 4.5. Model training and evaluation

The ADNI cohort was used for model training, while the NACC cohort served as an independent validation dataset. Within ADNI, participants with follow-up visits were randomly split into training and test sets, with 80% (n = 551) assigned to training. The remaining 20% (n = 154) and those with only single visit were reserved for testing. Multiple visits from the same participant were assigned to one group to avoid data leakage. Only training data were used to fit all components of the trajectory pipeline, including covariate adjustment, LNE model training, and PCA embedding. The trained models were then applied to the test set and external validation set to evaluate generalization. As the harmonized NACC dataset contains only single visit amyloid PET, it was used exclusively for external validation.

Model performance was evaluated using three complementary criteria. First, diagnostic classification accuracy was assessed by measuring how well branch assignment aligned with clinical diagnosis, specifically the separation between CN and AD participants based on their location on the trajectory branches. Given the potential heterogeneity of CN participants, we focus on those with Centiloid <10. Second, temporal consistency of the learned pseudotime was quantified using the violation ratio, defined as the proportion of consecutive visit pairs in which pseudotime decreases over time; lower values indicate better alignment with expected disease progression. Third, branch stability was evaluated by computing the proportion of longitudinal participants whose branch assignment changed across visits, reflecting the consistency of trajectory-based classification over time. Together, these metrics assess the biological validity, temporal coherence, and robustness of the learned trajectory.

### 4.6. Projection onto Trajectory

To project previously unseen individuals onto the 2D amyloid trajectory, their regional SUVR will first go through pre-trained linear adjustment model, then the pre-trained encoder in longitudinal embedding and finally the pre-trained PCA transformation to be located on the 2D trajectory. Their branch specific pseudotime will be obtained through the pre0trained spline function.

### 4.7. Association of branch-specific staging score with cognition

We examined the association between branch-specific staging score (i.e., pseudotime) and cognitive function to assess the clinical relevance of the inferred diverging branches. Cognitive performance was quantified using the Memory Score (PHC_MEM) and Executive Function Score (PHC_EXF)^32,41^. Cross-sectionally, Pearson correlation coefficients (r) and corresponding ρ values were calculated between branch-specific pseudotime and cognitive measures. These analyses were grouped by the identified trajectory branches to characterize branch-specific patterns of cognitive decline.

In addition, we also examine the longitudinal cognitive decline for CN/MCI participants located on two branches. To evaluate longitudinal cognitive trajectories stratified by branch assignment, linear mixed-effects models (LMMs) were fitted separately for memory (PHC_MEM) and executive function (PHC_EXF) scores in the CN plus MCI population of the ADNI cohort. The model took the form:

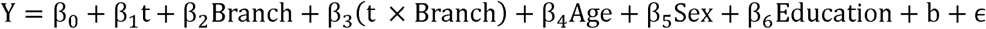

where Y is the cognitive score, t is follow-up time in months, and Branch is a binary indicator for AD-branch membership. The interaction coefficient β_3_ is the primary parameter of interest, capturing the difference in the rate of cognitive change between branches. A subject-level random intercept ∼ N(0, σ^2^_b_) accounted for individual baseline differences and the non-independence of repeated observations, and ∼ N(0, σ^2^) denotes the residual error.

Models were estimated using restricted maximum likelihood (REML) via the lme4 package in R^42^, and p-values for fixed effects were obtained using Satterthwaite’s degrees-of-freedom approximation as implemented in lmerTest^43^. The primary inferential quantity was the p-value for the time × branch interaction term, reflecting whether the two branches exhibited significantly divergent rates of cognitive change over follow-up. To ensure out-of-sample validity, analyses were conducted independently in the training and test splits of ADNI, and marginal predicted trajectories with 95% confidence intervals were derived from the fitted models using the ggeffects package^44^.

### 4.8. Survival analysis to assess the branch-specific AD conversion risk

To assess whether branch assignment predicted the risk of clinical progression to Alzheimer’s disease dementia, time-to-event analyses were performed using the Kaplan–Meier estimator and the Cox proportional hazards model^45,46^. Each subject was followed from their baseline cluster-assignment visit until first recorded conversion to an AD diagnosis (event) or their last available visit (censored), with time measured in months. Participants with a baseline label of CN and MCI who subsequently transitioned to AD within the longitudinal cluster trajectory were classified as converters. Survival curves were estimated separately for the AD-branch and normal-branch groups, and between-group differences were evaluated with the log-rank test. The magnitude of the association between branch and conversion risk was quantified by the hazard ratio (HR) and its 95% confidence interval from a univariable Cox model. All analyses were implemented using the survival^47^ and survminer^48^ packages in R and were replicated in both the training and test splits of ADNI to assess reproducibility.

### 4.9. Genome wide association analysis across branches

To investigate the genetic basis of trajectory-defined heterogeneity, we performed genome-wide association analyses (GWAS) to identify single nucleotide polymorphisms (SNPs) associated with branch membership. Branch assignment (AD-related vs aging branches) was treated as a binary outcome, and association testing was conducted using logistic regression under an additive genetic model implemented in PLINK 2.0^49^. Analyses included age and sex as covariates. In addition to the full cohort analysis, we performed a secondary GWAS restricted to cognitively normal (CN) and mild cognitive impairment (MCI) participants to identify genetic variants associated with divergence into AD-related versus aging trajectories at early disease stages. Genome-wide significance was defined at p<5×10^-8^

## Data availability

Harmonized amyloid PET data analyzed in this study were obtained respectively from the Alzheimer’s Disease Neuroimaging Initiative (ADNI) data portal (https://ida.loni.usc.edu) and the National Alzheimer’s Coordinating Center (NACC) through the Alzheimer’s Disease Sequencing Project (ADSP) Phenotype Harmonization Consortium. Access to these datasets is subject to the approval by the corresponding data repositories.

## Code availability

Pre-trained model together with best parameters is shared through GitHub (https://github.com/MZhao-ouo/LNE_Branch).

## Acknowledgements

Data collection and sharing for this project was funded by the Alzheimer’s Disease Neuroimaging Initiative (ADNI) (National Institutes of Health Grant U01 AG024904) and DOD ADNI (Department of Defense award number W81XWH-12-2-0012). ADNI is funded by the National Institute on Aging, the National Institute of Biomedical Imaging and Bioengineering, and through generous contributions from the following: AbbVie, Alzheimer’s Association; Alzheimer’s Drug Discovery Foundation; Araclon Biotech; BioClinica, Inc.; Biogen; Bristol-Myers Squibb Company; CereSpir, Inc.; Cogstate; Eisai Inc.; Elan Pharmaceuticals, Inc.; Eli Lilly and Company; EuroImmun; F. Hoffmann-La Roche Ltd and its affiliated company Genentech, Inc.; Fujirebio; GE Healthcare; IXICO Ltd.; Janssen Alzheimer Immunotherapy Research & Development, LLC.; Johnson & Johnson Pharmaceutical Research & Development LLC.; Lumosity; Lundbeck; Merck & Co., Inc.; Meso Scale Diagnostics, LLC.; NeuroRx Research; Neurotrack Technologies; Novartis Pharmaceuticals Corporation; Pfizer Inc.; Piramal Imaging; Servier; Takeda Pharmaceutical Company; and Transition Therapeutics. The Canadian Institutes of Health Research is providing funds to support ADNI clinical sites in Canada. Private sector contributions are facilitated by the Foundation for the National Institutes of Health (www.fnih.org). The grantee organization is the Northern California Institute for Research and Education, and the study is coordinated by the Alzheimer’s Therapeutic Research Institute at the University of Southern California. ADNI data are disseminated by the Laboratory for Neuro Imaging at the University of Southern California.

---

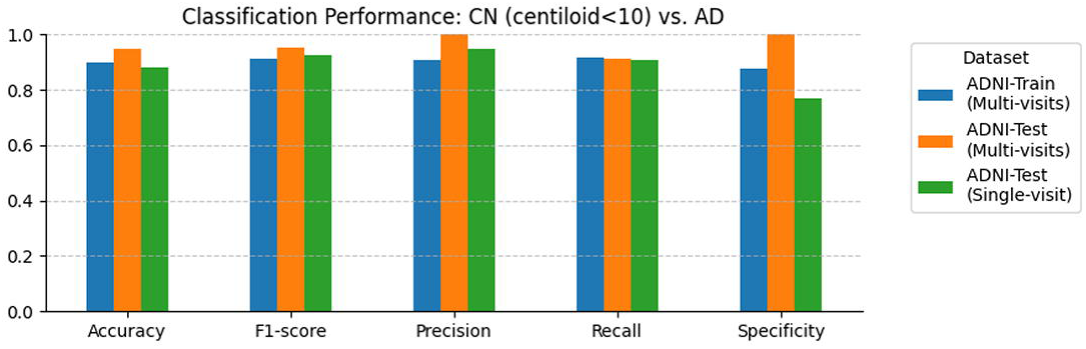

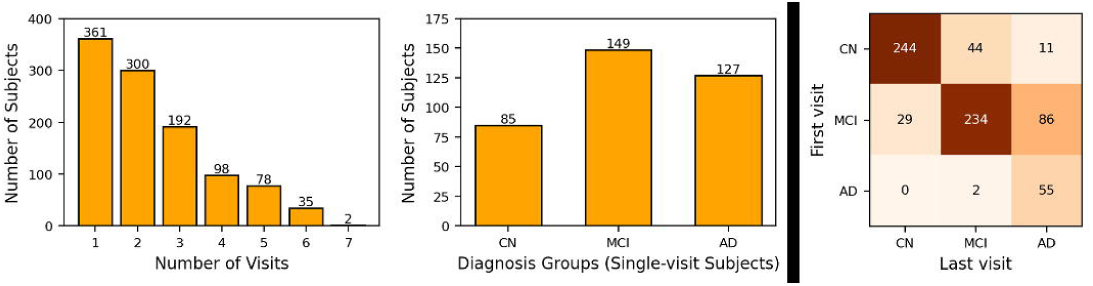

