## Supplemental Figure S1, Figure S2 and Table S1, Table S2, for "Divergent amyloid trajectories distinguish normal aging from Alzheimer’s disease progression"

### Supplementary Information

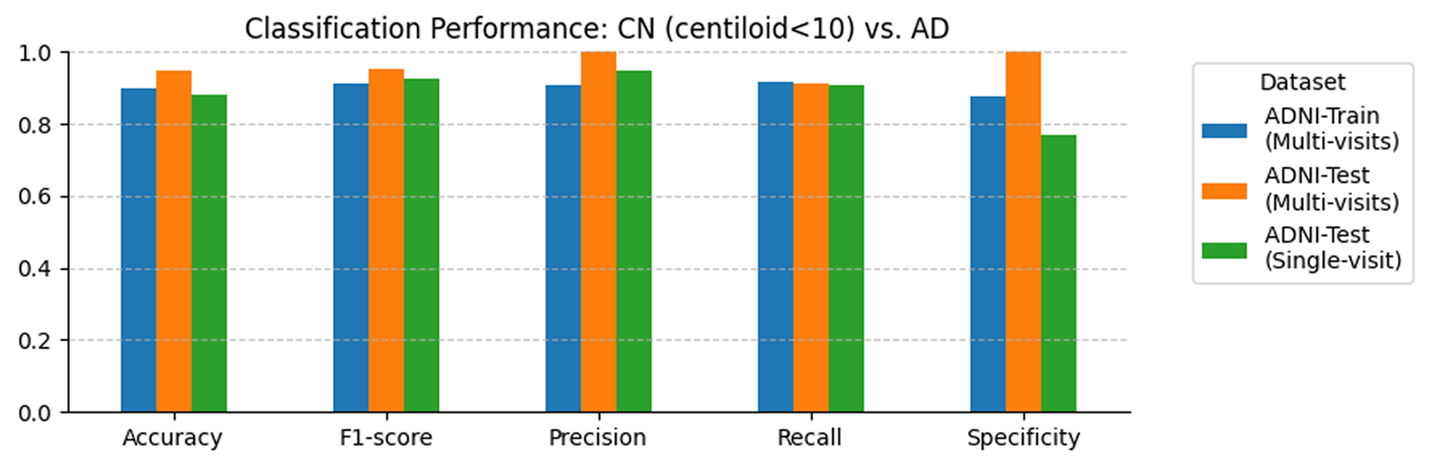

**Fig. S1: Branch-based classification performance for differentiating AD from amyloid-negative CN individuals (Centiloid <10).**

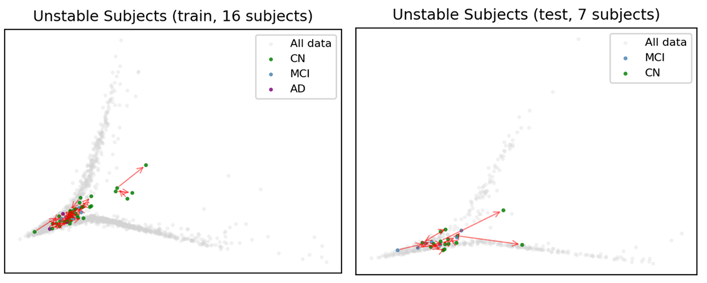

**Fig. S2: Branch switching among longitudinal subjects.** Subjects exhibiting branch switching across longitudinal visits in the training (left; *n* = 16) and test (right; *n* = 7) sets. Gray points denote all observations defining the trajectory, and colored points denote visits from branch-switching subjects, colored by clinical diagnosis. Red arrows indicate the temporal direction between longitudinal visits.

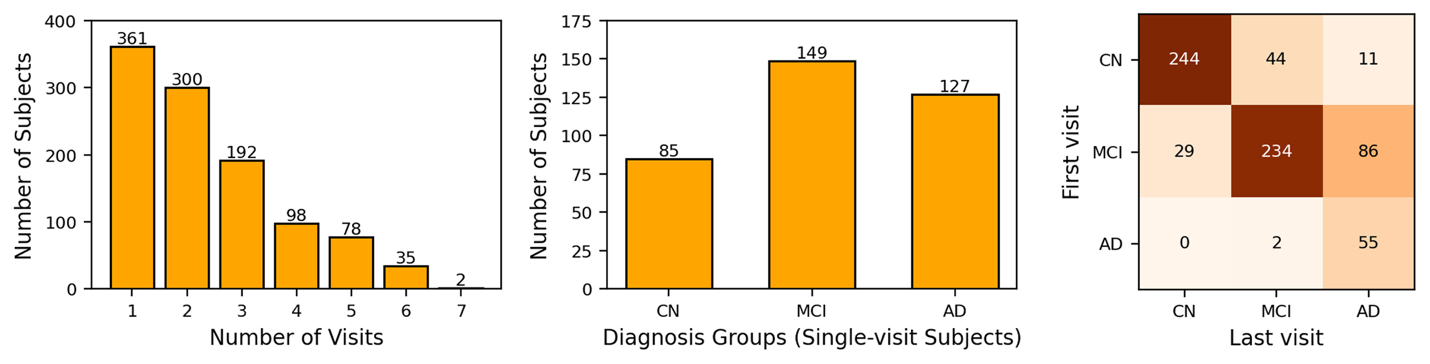

**Fig. S3: Overview of subjects from the ADNI cohorts.** (a) Distribution of the number of follow-up visits across all participants. (b) Diagnostic composition of participants with only a baseline visit. (c) Diagnostic composition of participants with multiple follow-up visits. Heatmap illustrates diagnostic transitions from baseline to the last available visit, where color intensity represents the proportion of subjects undergoing each transition.

**Table S1.** **CN & AD Classification performance of ADNI dataset**

|  | **Accuracy** | **F1-score** | **Precision** | **Recall** | **Specificity** |
| --- | --- | --- | --- | --- | --- |
| **Train** | 0.8989 | 0.9124 | 0.9083 | 0.9167 | 0.8750 |
| **Test**  **(multi-visit)** | 0.9464 | 0.9538 | 1.0000 | 0.9118 | 1.0000 |
| **Test**  **(single visit)** | 0.8828 | 0.9270 | 0.9474 | 0.9076 | 0.7692 |

**Table S2. Demographic information of different datasets (baseline)**

|  |  | **CN** | **MCI** | **AD** |
| --- | --- | --- | --- | --- |
| **ADNI** | **N** | 384 | 498 | 184 |
|  | **Sex**,  male/female | 179/205 | 294/204 | 109/75 |
|  | **Age**,  mean (SD) | 74.20 (6.72) | 73.16 (7.83) | 75.40 (7.68) |
|  | **Years of education**,  mean (SD) | 16.64 (2.57) | 16.11 (2.72) | 15.82 (2.63) |
| **NACC** | **N** | 260 | 12 | 49 |
|  | **Sex**,  male/female | 98/162 | 6/6 | 33/16 |
|  | **Age**,  mean (SD) | 72.81 (6.62) | 73.47 (9.75) | 73.64 (9.01) |
|  | **Years of education**,  mean (SD) | 16.63 (2.51) | 16.58 (2.78) | 21.00 (16.43) |
